# PeakBot: Machine learning based chromatographic peak picking

**DOI:** 10.1101/2021.10.11.463887

**Authors:** Christoph Bueschl, Maria Doppler, Elisabeth Varga, Bernhard Seidl, Mira Flasch, Benedikt Warth, Juergen Zanghellini

## Abstract

**Motivation:** Chromatographic peak picking is among the first steps in data processing workflows of raw LC-HRMS datasets in untargeted metabolomics applications. Its performance is crucial for the holistic detection of all metabolic features as well as their relative quantification for statistical analysis and metabolite identification. Random noise, non-baseline separated compounds and unspecific background signals complicate this task.

**Results:** A machine-learning framework entitled PeakBot was developed for detecting chromatographic peaks in LC-HRMS profile-mode data. It first detects all local signal maxima in a chromatogram, which are then extracted as super-sampled standardized areas (retention-time vs. *m/z*). These are subsequently inspected by a custom-trained convolutional neural network that forms the basis of PeakBot’s architecture. The model reports if the respective local maximum is the apex of a chromatographic peak or not as well as its peak center and bounding box.

In training and independent validation datasets used for development, PeakBot achieved a high performance with respect to discriminating between chromatographic peaks and background signals (accuracy of 0.99). For training the machine-learning model a minimum of 100 reference features are needed to learn their characteristics to achieve high-quality peak-picking results for detecting such chromatographic peaks in an untargeted fashion.

PeakBot is implemented in python (3.8) and uses the TensorFlow (2.5.0) package for machine-learning related tasks. It has been tested on Linux and Windows OSs.

**Availability:** The package is available free of charge for non-commercial use (CC BY-NC-SA). It is available at https://github.com/christophuv/PeakBot.

**Supplementary information:** Supplementary data are available at Bioinformatics online.

## 1 Introduction

Untargeted metabolomics approaches have gained much popularity in recent years. They aim at holistically detecting all compounds present in samples regardless of their chemical identity. Subsequently, statistical analysis is carried out to select differently abundant compounds between treatment and control conditions for further biological interpretation (Fiehn 2002).

The most used analytical technologies for detecting metabolites in aqueous samples are liquid or gas chromatography (LC or GC) coupled with high resolution mass spectrometry (HRMS) and nuclear magnetic resonance (NMR) spectroscopy. LC-HRMS offers the most versatility since it can be customized towards the samples or certain constituents of interest (e.g., polar or non-polar compounds, lipids, or secondary metabolites). GC-MS approaches are mostly used to study volatile compounds and NMR approaches offer the possibility for absolute quantification as it is not prone to matrix effects (Segers et al. 2019).

While in targeted approaches the substances to be investigated are defined by the experimenters and are typically available as authentic reference standards, in untargeted approaches novel compounds are also of particular interest. Depending on the metabolic capabilities of the organism under study, a high number of metabolites can be expected (Peisl, Schymanski, and Wilmes 2018; Alseekh and Fernie 2018). This high number of metabolic features combined with multiple experimental conditions and replicates makes it difficult to manually examine the raw-data and integrate each peak. Thus, automated software tools, which are capable of a) detecting chromatographic peaks in the dataset and b) integrate them reliably, are of utmost importance. These two steps are typically carried out simultaneously yielding an average mass-to-charge-ratio (*m/z*) and retention-time as well as a relative abundance (peak area or intensity of most abundant signal contributing to the peak).

The metabolomics community has recognized the need for automation early on and started developing holistic and reliable (open source) software approaches. Arguably, the best known approaches are XCMS (Tautenhahn, Böttcher, and Neumann 2008) and XCMS-Online (Tautenhahn et al. 2012), MzMine2 (Pluskal et al. 2010), MS-Dial (Tsugawa et al. 2020), MetAlign (Lommen and Kools 2012), OpenMS (Röst et al. 2016), and Lipid Data Analyzer (Hartler et al. 2017), among others; see also (O’Shea and Misra 2020). These tools scale well to different chromatographic conditions and MS instruments making them reliable and indispensable. However, as LC-HRMS methods can be customized (e.g., different chromatography methods, mass ranges, resolution, separation power, etc.), one software or parameter set does not fit all experiments. Furthermore, optimizing a data processing software for a particular analytical technique can be a challenging process that requires both an in-depth knowledge of the analytics as well as the data processing algorithm to assess the performance of different parameter sets.

Machine-learning has recently gained attention for the task of peak-picking with for example, NeatMS (Gloaguen, Kirwan, and Beule 2020). It does not perform peak-picking itself, but classifies peaks detected with other tools and rates the quality of the detected peaks (high quality, acceptable, noise). Thus, it can easily be integrated into existing workflows. Another machine-learning approach is peakOnly (Melnikov, Tsentalovich, and Yanshole 2020), which performs both classification and peak border estimation with the help of deep-learning. Both approaches utilize centroid data mode.

Here we seek to explore the possibility of using machine-learning convolutional neural networks (CNN) models for the task of detecting chromatographic peaks in single profile-mode chromatograms. In contrast to most algorithmic approaches (e.g., XCMS, MS-Dial, etc.), which rely on the optimization of method parameters, the presented method uses a series of user-provided references (i.e., chromatographic peaks and different background types) to custom-train a machine-learning model that can recognize such chromatographic peaks and calculate their respective analytical properties directly in the raw profile-mode data.

## 2 Approach

PeakBot is a python package that imports LC-HRMS datasets, pre-processes them and returns a list of detected chromatographic peaks. Figure 1 illustrates PeakBot’s processing pipeline. Unlike most available tools for chromatographic peak picking, the PeakBot framework uses profile-mode data instead of centroided data. Moreover, the presented approach does not have a region-of-interest (ROI) generation step as for example XCMS, but rather enumerates all local maxima (or a reduced subset), exports areas around this local maxima as standardized, super-sampled areas of the chromatogram and tests these areas for the presence of a chromatographic peak in its center. If the CNN model (Figure 2) detects a chromatographic peak, PeakBot also calculates the peak’s center and bounding box (retention-time and *m/z* dimension). PeakBot’s CNN model can also be trained for other categories of local maxima such as “walls” that span rather large parts of the chromatogram with only minimal random changes in intensity; see Supplementary Figure 1.

**Figure 1.**
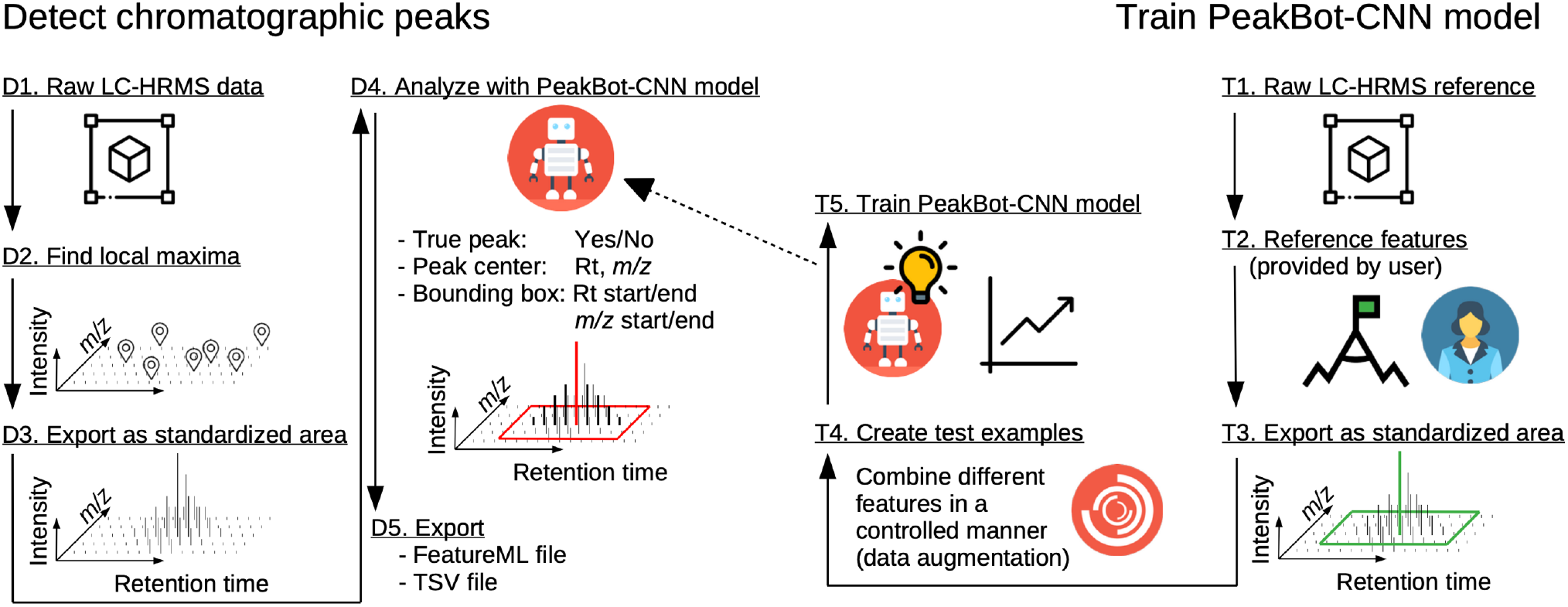
Overview of the detection and training steps of PeakBot. The left panel shows the steps for detecting chromatographic peaks with an already trained PeakBot model, while the right panel shows the steps of training a new PeakBot model from reference LC-HRMS data.

**Figure 2.**
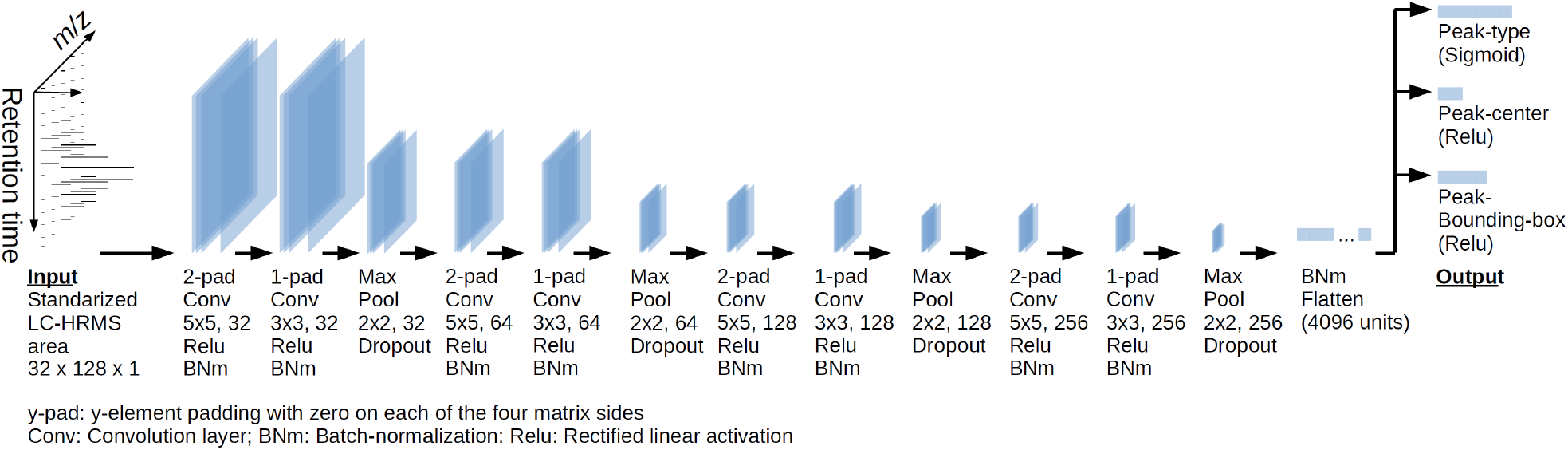
Overview of PeakBot’s CNN model. In its standard configuration PeakBot starts from the standardized LC-HRMS area and applies a series of convolution and max-pooling steps to the previous layer. Finally, the output of these convolutions is flattened and the type of the local maxima as well as the peak’s center and bounding box are determined.

### 2.1 Data processing

The main steps of detecting chromatographic peaks with PeakBot are illustrated in Figure 1 in the left panel. First, the profile-mode LC-HRMS data is loaded from files in the mzXML/mzML format. Then, all local maxima (i.e., individual *m/z* signals that are more abundant than their 8-neighboring *m/z* signals (1 scan earlier/later in the chromatographic domain and preceding/succeeding *m/z* signal with an offset less than the median difference between typical neighboring *m/z* signals of a profile mode peak), Moore neighborhood) are detected. At this stage the detection is carried out regardless of whether the apex signals represent a true chromatographic peak or not.

All detected local maxima are exported as standardized LC-HRMS areas consisting of a pre-set number of HRMS scans (default 32) and a pre-set number of *m/z* signals (default 128). These pre-set defaults can be increased for example for chromatographic peaks with a high number of data points (it is recommended that the size of HRMS scans in the standardized area should be at least 2-3 times the average number of scans that map to a typical chromatographic peak). Because on some HRMS instruments slight variations in the accuracy of the *m/z* signals can occur, even on adjacent MS scans, these standardized areas are super-sampled. This means that an exact, equally spaced number of reference *m/z* values relative to the respective local maximum is generated with the local maxima being in the center of this area. Finally, the standardized area is scaled to a maximum abundance of value one.

As the number of local maxima in an LC-HRMS sample can be quite high (e.g., due to high-frequency noise), an optional pre-processing step of PeakBot is a basic gradient descend-based peak detection approach. This pre-processing starts from each detected local maximum. Adaptive EICs are calculated for each local maximum (see Supplementary Figure 2). A fixed *m/z* value is not used but rather the EIC adapts from the previous *m/z* value of the previous scan thereby accounting for small shifts between the profile-mode signals. Each EIC is smoothed with a Savitzky-Golay filter. Starting from each local maximum the pre-processing algorithm moves left and right on the retention-time axis until the inflection points are reached (strictly monotony decreasing EIC). Then each EIC is further traveled until either a scan has an increase in the intensity relative to its predecessor or until the intensity relative to the peaks’ apex (the local maximum) undercuts 1%. If this pre-processing step is used, then only local maxima with a minimum peak width are further used and exported, while the others are prematurely discarded.

Each standardized area is used as the input for the CNN model, which tests it for the presence of a chromatographic peak in its center. PeakBot reports detected chromatographic peaks either as solitary peaks or such having left and/or right isomeric neighboring peaks (partly overlapping chromatographic peaks). If the local maximum and thus the standardized area does not contain a chromatographic peak, but for example background walls, no peak will be reported but rather the category wall or background. For local maxima designated to contain chromatographic peaks, PeakBot will also calculate the peaks’ center (which will mostly be the center of the standardized area) as well as a bounding-box in both the retention-time and *m/z* dimension.

Chromatographic peaks areas are calculated by summing up all signal intensities that are within the reported boundaries. This also applies to partly-co-eluting or shoulder peaks, as the boundaries of such peaks are automatically updated during the training instance generation (see next section).

Finally, detected chromatographic peaks are exported to a tsv-file and as a featureML file that can be used to visualize the results e.g. with TOPPView (Sturm and Kohlbacher 2009).

### 2.2 Training of PeakBot

The main steps of training a new PeakBot-CNN model are illustrated in Figure 1. To train a new PeakBot model from scratch, reference LC-HRMS chromatograms and a list of reference features must be provided by the user. The reference chromatograms do not need any special characteristics, however the list of reference features provided by the user must only consist of features that represent single, isolated ions with as little background as possible. Partly co-eluting isobaric or isomeric compounds are not supported yet and cannot be used as reference features. In a typical untargeted metabolomics experiment compiling such a list should be easily with standard compounds, QC samples and even unknown metabolites. This list of reference features can also be extended easily by using different isotopologs of already available compounds determined by the user. For each reference feature the user must specify the peak center (retention-time and *m/z* value) and its bounding box in the retention-time and *m/z* dimension. Furthermore, the user must specify a couple of background-regions that contain backgrounds.

Once specified, PeakBot first recognizes the features in the different samples and calculates their properties directly from the data (weighted-average *m/z* and apex retention-time) thereby accounting for small shifts and drifts. For this refinement step the basic gradient-descend pre-processing algorithm is utilized.

PeakBot uses this user-curated input to generate many diverse instances and are used as the input for training a new PeakBot-CNN model. Specific characteristics of the LC-HRMS dataset and especially those of the references provided by the user (narrower/broader chromatographic peaks, deviation of related *m/z* signals, saturation effects, etc.) are retained in the automatically generated training instances. Each such instance has a ground-truth center representing the local-maximum. This ground-truth can be either a feature or a background and the respective area in the chromatogram is super-sampled identically to the super-sampling carried out during the detection phase of PeakBot. To avoid over-fitting and to automatically generate many training instances (data augmentation), this ground-truth and super-sampled area are iteratively combined with randomly selected other features/background signals (decoys) from the user’s input thereby simulating a diverse neighborhood of the local maximum. These other features/background signals are randomly placed into the standardized area of the ground-truth. A variable degree of overlap (maximum 50% on the peak width and maximum 33% on the peak-area intersection over union IOU) is allowed to simulate non-baseline separated chromatographic peaks. The peaks’ boundaries are automatically updated when another peak is placed next to a chromatographic peak as these two form a partly co-eluting or shoulder peak. During training the numbers of these different classes of training instances (i.e., baseline separated, overlapping peaks and backgrounds) are tried to be kept similar. Additionally, all signals are randomly varied (multiplied with a random value, default within the range of 0.9 to 1.1) to simulate intensity-deviations caused by the instrument.

Each such generated and augmented training instance is then used by PeakBot to optimize the CNN model with the aim of correctly detecting the training instance, classify it and calculate its properties.

### 2.3 PeakBot-CNN model

PeakBot utilizes a convolutional neural network (CNN) that has a standardized LC-HRMS area as input. Then, several convolutional steps and pooling layers follow. Two convolutional layers are followed by a max-pooling layer and a ReLu activation. This cascade is repeated four times each time reducing the intermediate size but increasing the number of computed information gained from the layers. The number and computed information can easily be extended if required by the user. After the last convolutional layer, the convoluted area is flattened, from which the outputs (feature-type, boundaries, and center) are calculated. These outputs are A) peak type (one-hot-encoded peak category consisting of 6 types in its standard configuration), B) peak-center (indices in the retention-time and *m/z* dimension in the standardized area), and C) peak-bounding-box (start/stop indices in the retention-time and *m/z* dimension in the standardized area). The CNN architecture is illustrated in Figure 2.

Training is performed with the TensorFlow framework. For optimizing the output of the model, different loss functions are applied for the local-maximum types (loss: categorical-crossentropy), peak-center (loss: Huber), peak-bounding-box (loss: Huber), peak mask (loss: binary crossentropy). Moreover, also the peak accuracy (loss: categorical accuracy) and the peak-bounding-box intersect over union (loss: custom) are reported to the user but these two are not used to optimize the model.

### 2.4 Grouping of results from different chromatograms

To group the detected chromatographic peaks from different LC-HRMS chromatograms, PeakBot implements a k-nearest-neighbor approach with a user-defined retention-time and *m/z* window. Further details about this algorithm are included in Supplementary Information 1.

This grouping step can be repeated multiple times with wider and narrower parameters thereby iteratively aligning the chromatograms to each other. If experimental conditions with vastly different metabolic constituents are analyzed, the different experimental groups can be aligned independently first to create a virtual consensus sample and then the results of different experimental conditions can be merged.

Finally, a comprehensive data matrix is generated consisting of all detected chromatographic peaks and their abundances in the different samples. If a chromatographic peak has not been detected in a sample, the respective cell will be empty (missing value). Furthermore, the results can be exported to a featureML file for visualization with e.g. with TOPPView (Sturm and Kohlbacher 2009).

### 2.5 Implementation details

PeakBot is implemented in the python programming language (at least 3.8). The machine-learning CNN model is implemented with the tensorflow package (at least 2.5.0, https://www.tensorflow.org/).

PeakBot utilizes the numba package (http://numba.pydata.org/) for GPU compilation to detect local maxima in the LC-HRMS data and export them to the standardized areas. The generation of the augmented training instances also benefits from the numba package as this functionality can also be executed on CUDA-enabled graphics cards. Alternatively, a CPU version with just in time compilation of python code is available.

PeakBot reads the mzXML/mzML file format for raw LC-HRMS data.

## 3 Methods

### 3.1 Training and validation datasets

To train a new CNN model and monitor PeakBot’s performance during the training step, different datasets were generated from the user-provided reference feature list. Briefly, these were derived from untargeted metabolomics experiments of wheat ears (named “WheatEar”, further information is provided in Supplementary Information 2), porcine hepatic microsome incubation samples (named “PHM”, further information is provided in Supplementary Information 3) and 4 other datasets obtained from the MetaboLights (https://www.ebi.ac.uk/metabolights; MTBLS1358 (Flasch et al. 2020), MTBLS868 (Mbekeani et al. 2019), MTBLS797 (Raheem et al. 2019)) and the Metabolomics Workbench (https://www.metabolomicsworkbench.org/; ST001450 (Zhang, Dong, and Raftery 2020)) repository (further information about these datasets is provided in Supplementary Information 4). Separate PeakBot CNN models were trained for each dataset to evaluate the performance as presented in Section 2.2. Cross validation (5-fold) with randomly selected peaks for training and validation was used.

For each such LC-HRMS dataset, four reference-set were generated for training and verifying the CNN models’ performance. These sets are:

T: The training set obtained from the training chromatograms with a list of at least 100 reference features.

V: A validation set obtained from the training chromatograms (same as for T) but different reference features (no overlap with T).

iT: A validation set obtained from different validation chromatograms (no overlap with T), but the reference features used for training the model (same as in T).

iV: A validation set obtained from different validation chromatograms (no overlap with T) and also different reference features (no overlap with T). As this set neither shares the chromatograms nor the reference features with the training set T it can be considered a<u>n</u> independent validation set of the model.

During training, the agreement of the ground truth (i.e., the generated training instances) and the predictions of the model are summarized in several loss/metric values detailed in Table 1. All validation results were critically evaluated with respect to the metrics reported by the training process. Moreover, a subset of all results was also manually verified by the person training the CNN model.

**Table 1.**
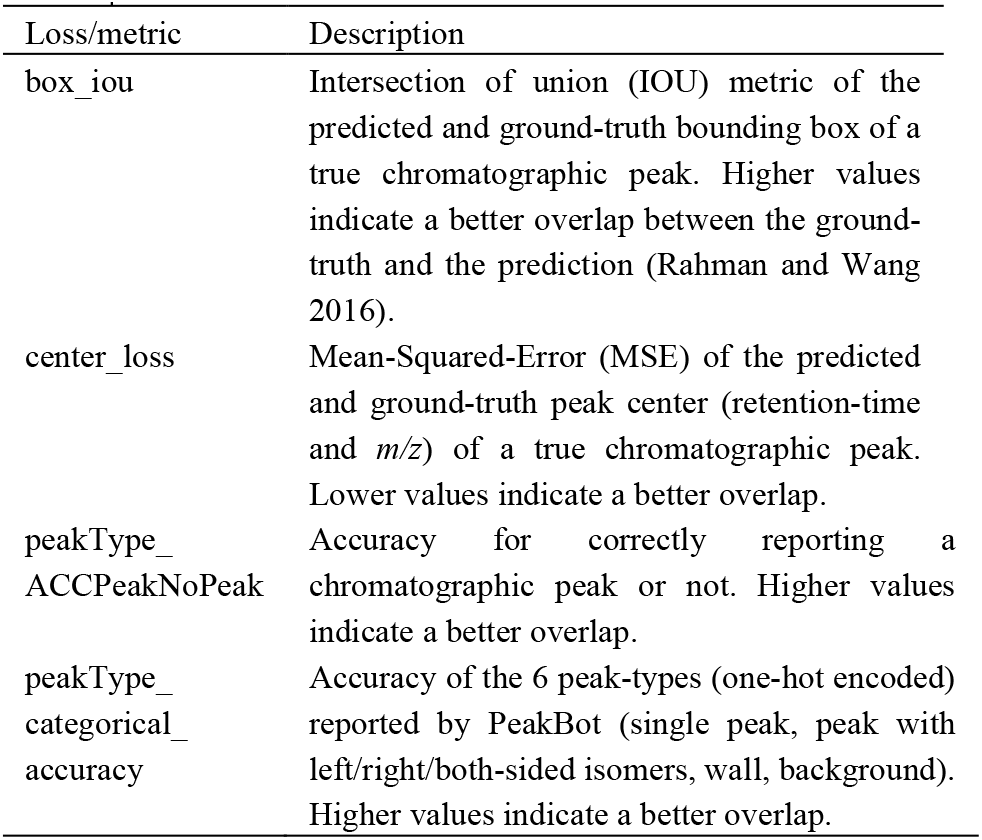
Overview of loss/metric values.

| Loss/metric | Description |
| --- | --- |
| box_iou | Intersection of union (IOU) metric of the predicted and ground-truth bounding box of a true chromatographic peak. Higher values indicate a better overlap between the ground-truth and the prediction (Rahman and Wang 2016). |
| center_loss | Mean-Squared-Error (MSE) of the predicted and ground-truth peak center (retention-time and $m/z$) of a true chromatographic peak. Lower values indicate a better overlap. |
| peakType_ACCPeakNoPeak | Accuracy for correctly reporting a chromatographic peak or not. Higher values indicate a better overlap. |
| peakType_categorical_accuracy | Accuracy of the 6 peak-types (one-hot encoded) reported by PeakBot (single peak, peak with left/right/both-sided isomers, wall, background). Higher values indicate a better overlap. |

## 4 Results and Discussion

Humans are exceptionally good at visually recognizing structure in unstructured data (e.g., object detection and separation, recognition of (co-eluting) chromatographic peaks). However, it is not possible for them to pick many peaks manually as humans are unfortunately very slow at this task. Thus, reliable, automated peak-picking is important.

Here PeakBot is presented. It is a CNN-machine learning model for the detection of chromatographic peaks in LC-HRMS datasets. As the analytical parameters of different datasets (e.g., chromatographic peak width and/or *m/z* deviation) can be quite different, the PeakBot CNN models can be custom-trained for each dataset or analytical method (e.g., scan rate for UHPLC or TOF instruments). The trained CNN model then inspects new chromatograms for the presence of chromatographic peaks or to different types of backgrounds observed in the dataset.

PeakBot has different outputs namely peak-type, -center and -bounding-box. The peak-type is a one-hot-encoded class identifier and consists of four types if a true chromatographic peak is believed to be in the center of the standardized area. These four peak types are whether the peak is isolated (no left and right isomeric neighbors) or not. Additionally, two peak categories also indicate if the local maximum is a background signal (either a wall or a random signal). If PeakBot considers the local maximum to be a true chromatographic peak, it will also calculate the peak’s center and bounding box, which describe the location of the peak in the two-dimensional chromatogram. An example of the detection of local maxima and classification as true chromatographic peaks and backgrounds is shown in Supplementary Table 1.

### 4.1 Training and validation of PeakBot

Training of machine-learning models is a difficult process with different aspects to be considered as otherwise the model might overfit. In this respect overfitting refers to a model that has a minimal error on the training dataset but does not generalize well with new data not used for the training process and thus has a poor prediction quality for new data. It is common practice to test the performance of machine-learning models during training also with independent validation instances (here iV) to check for such effects. If the model generalizes well, the validation- and training-reference-sets should have similar loss and metric values.

Three validation data sets are used to verify and judge the trained model’s performance. For this, at least one independent reference-set (i.e., same analytical method, but different features and chromatograms than used for training) are used. The losses and metrices for the training and validation for the six demonstration datasets are illustrated in Figure 3 and Supplementary Figure 3. The loss and metric values were similar (less than 5% difference in the metrices) between the training (T) and the three validation reference-sets (V, iT, iV) indicating that the model was not overfitted to the training data. The model was able to accurately estimate if a local maximum was indeed a chromatographic peak (accuracy for the peakType and binary classification for peak or no peak). In this respect, the independent reference-set iV was most interesting as it was derived from chromatograms and reference features completely independent of the training data. The trained PeakBot models were able to recognize chromatographic peaks or backgrounds with an accuracy of more than 99% (peakType_ACCPeakNoPeak metric) on average. The peaks’ centers were similar to the training and other validation reference-set (center_loss), while the bounding-box agreed a little less (box_iou).

**Figure 3.**
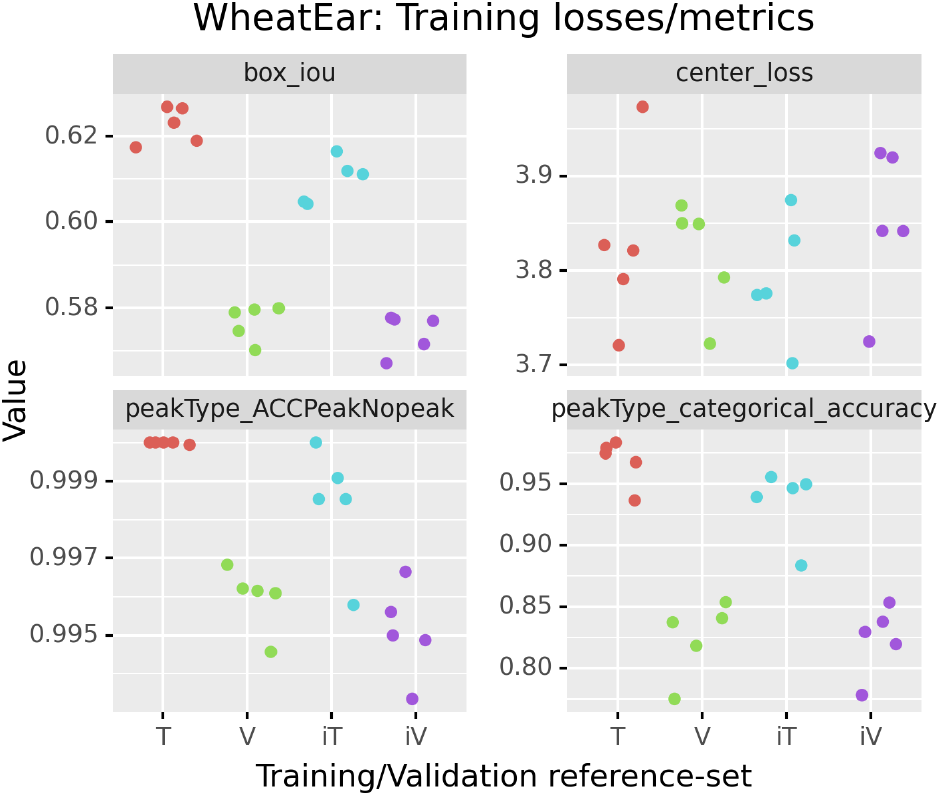
Overview of training losses and metrices on the training dataset T and the additional validation sets V, iV, iT.

### 4.2 Size and composition of reference feature set

PeakBot requires reference features for training of a CNN model. These reference features must be isolated chromatographic peaks curated by the user. Moreover, the different reference features should cover all expected, typical peak shapes of the analytical method.

To investigate the effect of differently sized reference lists upon the CNN-models performance, subsets of different size were generated from the initial reference feature list consisting of 2,731 reference features, which were combined to 1,048,576 training instances. Different sizes (10 to 2400) of the reference list were used to train separate PeakBot CNN models and to obtain their metric values.

In summary, a slight reduction in performance was observed when fewer reference features were used during the training phase (Supplementary Figures 4 and 5). A change in metric values was observed at around 100 features. Using a lower number of features greatly reduced the performance of the model, while the model did not noticeably benefit from a higher number of reference features. Nevertheless, the different models were still able to differentiate between true chromatographic peaks or backgrounds with a high certainty. Thus, this investigation showed that at least 100 reference features (including isotopolog features) should be provided during training by the user for good performance. Only the approximate retention-time and *m/z* values (i.e., within the actual peak’s boundaries) of the reference features need to be provided by the user as PeakBot will automatically adapt to the respective properties of the chromatographic peaks (apex in retention time and apex in *m/z* dimension) in the reference chromatograms.

Different sets of reference features were used for the training process to simulate researchers likely to assemble unequal lists of ground-truth data for training. The results are illustrated in Supplementary Figure 4 and 5. For a fixed training length, a replicate is a randomly selected set of the large training set initially available. Only minor differences in the performance metrics are reported for the different repetitions of the training indicating a good generalization of models’ detection process. Moreover, as only a certain number of the large instance set was used for training and the remaining features were used for verification in the iT, V and iV reference-set, it can also be concluded that PeakBot generalizes well to similar chromatographic peaks in the dataset. Furthermore, an already trained and validated model can easily be reused for processing datasets with similar chromatographic peak characteristics (i.e, data generated with the same analytical method as used for the initial training of the PeakBot model).

### 4.3 Comparison of PeakBot with XCMS, MS-Dial and peakOnly based peak-picking

To compare the performance of PeakBot with that of other peak-picking approaches, we compared it to XCMS, MS-Dial and peakOnly. XCMS and especially its centWave algorithm is a popular software package for detecting chromatographic peaks in LC-HRMS data in an untargeted manner (Tautenhahn, Böttcher, and Neumann 2008). It utilizes a wavelet-based algorithm to detect chromatographic peaks of different widths. The MS-Dial software is another software for LC-HRMS data processing (Tsugawa et al. 2020). It uses a gradient-descend inspired peak detection method and offers the user a sophisticated graphical user interface for easy data-processing. Finally, peakOnly is a machine-learning based peak-detection software, similar to PeakBot, and can be employed for peak-picking and comparison (Melnikov, Tsentalovich, and Yanshole 2020).

The comparison of these peak-picking algorithms was carried out on the PHM dataset. All four algorithms were independently optimized to the PHM dataset or a custom-dataset trained PeakBot CNN model. Then, all detected peaks were compared automatically to see how well these methods agree in respect to detecting the chromatographic peaks in an untargeted manner. It was necessary to allow for a large retention-time shift of 10 seconds and a large *m/z* shift of 10 ppm as the tools calculated the respective values differently. Supplementary Information 5 provides details of this automated comparison.

The results of this automated comparison for the PHM dataset are illustrated in Supplementary Figure 6. A high number of features (on average 73%), were successfully detected with all four approaches. On average another 12% of the features were not detected with peakOnly but with the remaining three tools and another 4.7% were detected exclusively with MS-Dial, peakOnly and PeakBot. The remaining combinations of two tools or each tool individually resulted in less than 2% additional features on average. This demonstrates that all tools did a good job at picking chromatographic peaks in general, although there are differences between the tools.

The results obtained for one chromatogram (sample 08_EB3391_AOH_p_60) were also compared manually using TOPPView (Sturm and Kohlbacher 2009) and are illustrated in Figure 4. A maximum of 50 features were used from each Venn-intersection. Of the 18 features detected only by PeakBot 78% were true chromatographic peaks. The features detected only by XCMS, MS-Dial and peakOnly were mostly incorrectly reported chromatographic peaks and actual background walls. Moreover, a high number of true chromatographic peaks were present in any Venn-intersection where PeakBot classified the respective features as true chromatographic peaks (red circle in Figure 4). The combination of XCMS and MS-Dial results also showed a high confidence in the chromatographic peaks detected only with these two approaches. Interestingly, 24 of these additional peaks were of high quality and abundance but with almost baseline separated isomeric other compounds. The remaining features (between 2 and 14) were detected by combinations of MS-Dial, XCMS, and peakOnly and represented mostly incorrectly classified backgrounds.

**Figure 4.**
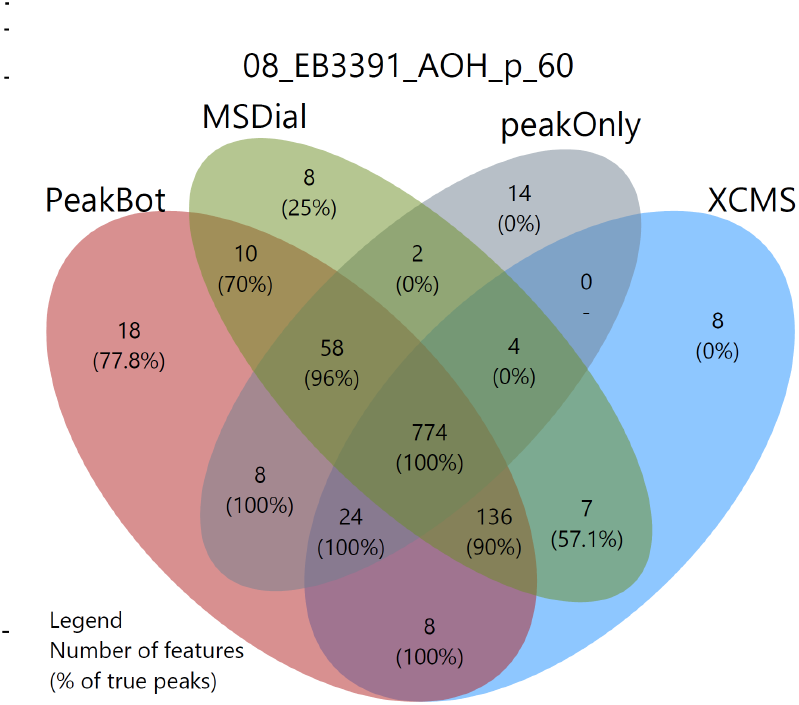
Comparison of results from PeakBot, XCMS, MS-Dial and peakOnly for a selected sample of the PHM dataset.

In summary this overview demonstrates that PeakBot performs better at detecting and classifying true chromatographic peaks. In these comparisons it detected on average 94.65% of all features detected with the different approaches and furthermore its classification accuracy was high, especially when the features were only detected solely with PeakBot or one additional data processing approach.

### 4.4 Relative quantification

Reliable peak detection and peak area integration for subsequent statistical comparison of different experimental conditions are the main tasks of peak picking. For optimal and reliable peak area integration, PeakBot automatically derives peak borders for detected chromatographic peaks. These borders are used for peak area integration. To test how well and repeatable the trained CNN model was able to integrate the peak areas, the data was a) manually sighted to confirm correctly detected peak areas, b) a comparison of the replicates from the WheatEar study using relative standard deviation (RSD) was carried out, and c) peak areas determined from PeakBot and XCMS were compared.

The manual, human verification confirmed that PeakBot reliably detected the peaks’ borders in the profile-mode data in both the retention-time and *m/z* dimension. In addition, for partly co-eluting isomeric peaks, PeakBot successfully separated these two chromatographic peaks and calculated their boundaries accordingly.

The RSD values of the replicates from the WheatEar study, which were technical replicates of repeated injections from the same sample at different injection volumes, showed small sample-to-sample differences including both instrumental and the data processing variations. On average less than 10% RSD was observed for the different features and 90% of them had RSD values lower than 20%, which was well within the typical ranges of untargeted metabolomics experiments (Bueschl et al. 2014). The distribution of the RSD values is illustrated in Figure 5A.

**Figure 5.**
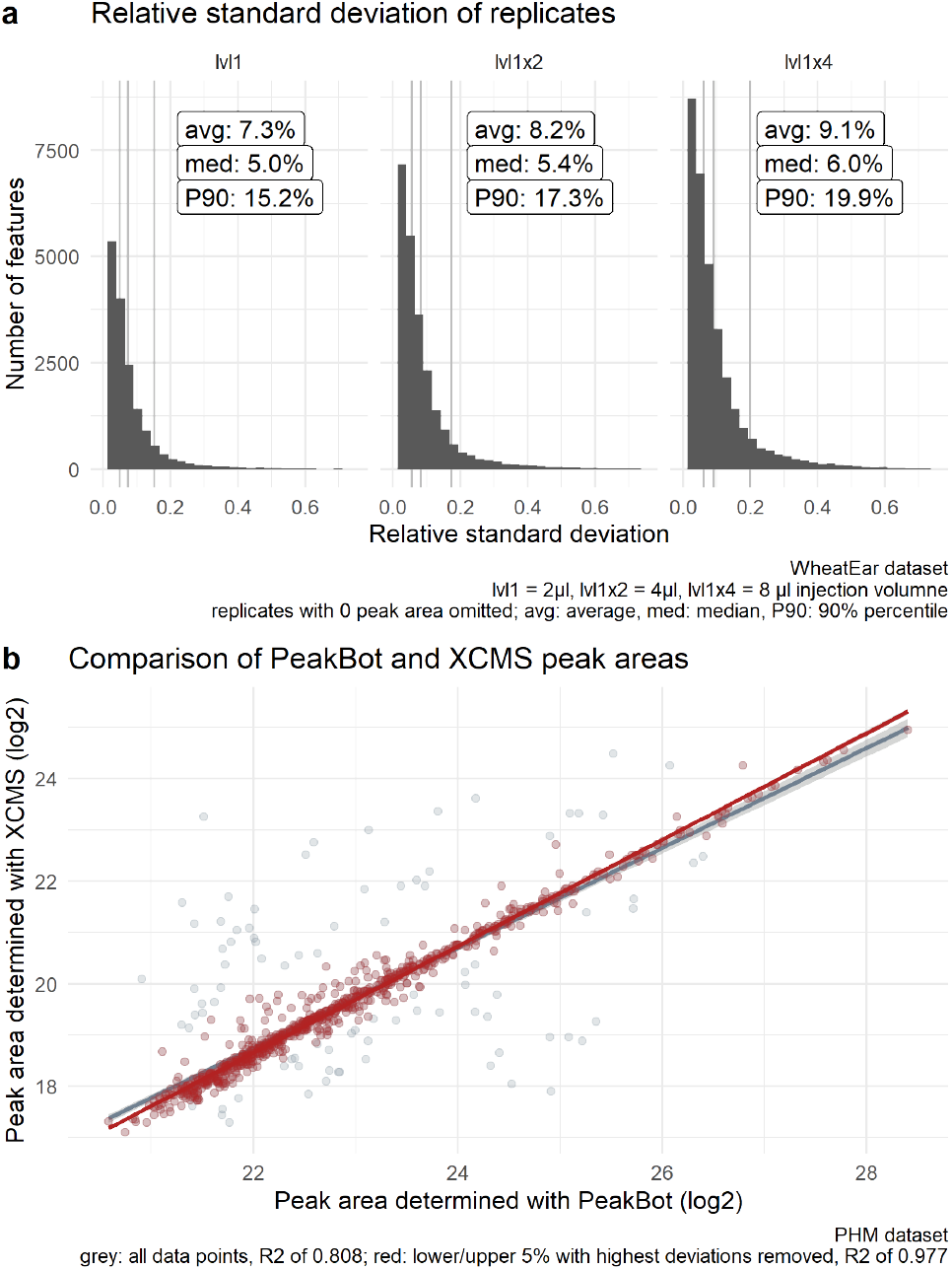
a RSD values of the peak areas of the WheatEar dataset. b Comparison of peak areas integrated with PeakBot and XCMS.

A comparison between the peak areas determined with PeakBot and XCMS from the tool comparison in the previous section showed a linear correlation of features detected with both approaches. When all features were taken into account, the R^2^ value was 0.808. This rather low R^2^ value is of no surprise, as outliers (i.e., incorrectly integrated peaks with either method) are expected with every peak-picking tool. When the bottom and top 5% of features with too dissimilar peak areas (i.e., 10% of the features in total) were omitted, the R^2^ value improved to 0.977. The linear relationship between PeakBot and XCMS is illustrated in Figure 5B. This comparison also showed that the peak areas determined with PeakBot were much higher than the peak areas determined with XCMS, but this was to be expected as PeakBot works with profile-mode data. The factor between the corresponding peak areas were on median 9.96 with the 25% and 75% percentile being 9.42 and 10.78.

These three investigations confirm that PeakBot accurately determines the peaks’ boundaries.

### 4.5 Runtime

CNN models are typically computationally more complex than the commonly utilized algorithmic approaches that work with one-dimensional EIC data. It is thus of no surprise that the data processing with PeakBot takes longer in comparison. However, PeakBot uses the TensorFlow framework for its machine-learning related tasks and thus it has the possibility to harness the computational power of graphical processing units (GPUs) of CUDA-enabled graphics cards. While such a setup does not allow processing several chromatograms in parallel, it allows processing many local maxima of a single chromatogram in parallel thereby reducing the overall computation time. As a result, it is recommended to run PeakBot on a PC equipped with a CUDA-enabled GPU. However, a CPU-only version that utilizes just-in-time compilation of python to C code with the numba package (https://numba.pydata.org) is also available, but processing times are much longer than for the GPU version.

All run-time tests were carried out using either an Nvidia Tesla V100S or an Nvidia GTX 970 GPU. The first one is currently one of the best performing CUDA cards available, however, it is typically installed only in super-computing centers, while the latter one is a some 6-year-old mainstream/gaming graphics card. On both systems the CNN-model and the necessary pre-processing routines that are also implemented with GPU support executed within reasonable time demonstrating that PeakBot’s pre-processing and CNN model execute quickly even on older, desktop-PC hardware. The systems and the runtimes of generating the training reference-set, training a new PeakBot model, as well as detecting chromatographic peaks in LC-HRMS samples are summarized in Table 2.

### 4.6 Generation of MSMS exclusion lists

Data dependent acquisition (DDA) of MS/MS spectra is typically carried out in untargeted metabolomics experiments to support feature annotation/identification. However, large backgrounds (i.e., walls) especially in samples with low metabolite ion abundances can interfere with this acquisition strategy and thus the MS/MS coverage of the sample’s metabolites is reduced. The degree of the reduction depends on the background and its intensity. An example of such a chromatogram where mainly parts of the background have been used for DDA is shown in Supplementary Figure 7.

In such experiments PeakBot’s background annotation feature can be useful for the automated generation of MS/MS exclusion lists for subsequent measurements. Such exclusion lists can be generated e.g., from the detected walls in background samples. In the shown example the use of such an exclusion list increased the MS/MS coverage of true chromatographic peaks increased from 1,356 to 1,605 in negative and 855 to 1,810 in positive ionization mode, respectively (Supplementary Table 2), which corresponds to an increase of 18% and 112% respectively. Nevertheless, it should be noted that peaks with the same *m/z* ratio as such background walls could potentially be missed with this strategy.

## Summary

PeakBot is a high-performance machine-learning based approach for detecting chromatographic peaks in LC-HRMS profile-mode datasets. The use of reference features and chromatograms makes it independent of different parameter settings and thus it can adapt to the dataset’s characteristics. Furthermore, the advantage is that PeakBot reports metrices about its training-progress thereby allowing the user to monitor its performance on an independent reference-set not used for training but for validation. To train a new model, PeakBot provides functions to automatically generate diverse training instances thereby reducing the number of reference features to be provided by the user.

For prediction, PeakBot requires only the raw chromatograms in profile-mode data and an intensity threshold. All local maxima (or alternatively these with a gradient-descend peak shape) are used as starting points for the detection of chromatographic peaks. Each local maximum is then inspected by the reference-feature trained model for chromatographic peaks or backgrounds.

In our tests, PeakBot showed to be a reliable peak-picking method and was able to compete with established software tools. Moreover, PeakBot also reliably integrates the peaks’ boundaries for relative quantification thereby not introducing a too high data processing variation.

Additionally, as PeakBot also classifies background signals it can be used for the efficient and reliable generation of exclusion lists for broad walls, which are often unnecessarily used for MS/MS scans thereby saving experimentalists time, resources, and sample.

PeakBot demonstrated to be a viable and good alternative for peak picking. As PeakBot requires real LC-HRMS data for its training, different models can be tuned for certain LC-HRMS characteristics (such as long fronting or tailing, zero/missing intensity values in the EICs, strong noise or non-Gaussian peak shapes). As a python package PeakBot also allows programmatic access to the raw LC-HRMS data and will be of interest for prototyping ideas. Furthermore, it can be integrated in other software tools and the models can easily be reused or shared. In future releases of the framework, we want to add machine-learning capabilities also to the tasks of grouping/aligning detected features from different results, re-integration, convolution, and annotation of features and provide an easy-to-use graphical user interface for users without coding experience.

PeakBot and related examples are available free of charge for academic use at https://github.com/christophuv/PeakBot and https://github.com/christophuv/PeakBot_example.

## Acknowledgements

We acknowledge Eszter Borsos for performing the porcine hepatic microsome assays, Lydia Woelflingseder for performing cell experiments, and Doris Marco and Rainer Schuhmacher for scientific input. The authors thank the members of the Mass Spectrometry Center at the University of Vienna. This project was supported by the BOKU Core Facility Bioactive Molecules: Screening and Analysis. We express our gratitude to the authors of the publicly available datasets we used from MetaboLights and Metabolomics Workbench. We also want to acknowledge and thank the anonymous reviewers, who provided us with critical feedback to improve the method and this manuscript. Figures have been designed using resources from https://flaticon.com.

