## Supplementary Materials for "PeakBot: Machine learning based chromatographic peak picking"

[1]Department of Analytical Chemistry, University of Vienna, Waehringer Str. 40, A-1090 Vienna, [2]IFA-Tulln, University of Natural Resources and Life Sciences Vienna, Konrad-Lorenz-Str. 20, A-3430 Tulln, [3]University of Natural Resources and Life Sciences, Core Facility Bioactive Molecules: Screening and Analysis, Konrad-Lorenz-Str. 20, A-3430 Tulln, [4]Department of Food Chemistry and Toxicology, University of Vienna, Waehringer Str. 38-40, A-1090 Vienna

**Supplementary information:** Supplementary data are available at *Bioinformatics* online.
**Note:** Source code and scripts for generating the results presented in the manuscript and this Supplementary Information is presented on https://github.com/christophuv/PeakBot_Publication and https://github.com/christophuv/PeakBot_example.

### Supplementary Figure 1

Illustration of a wall background.

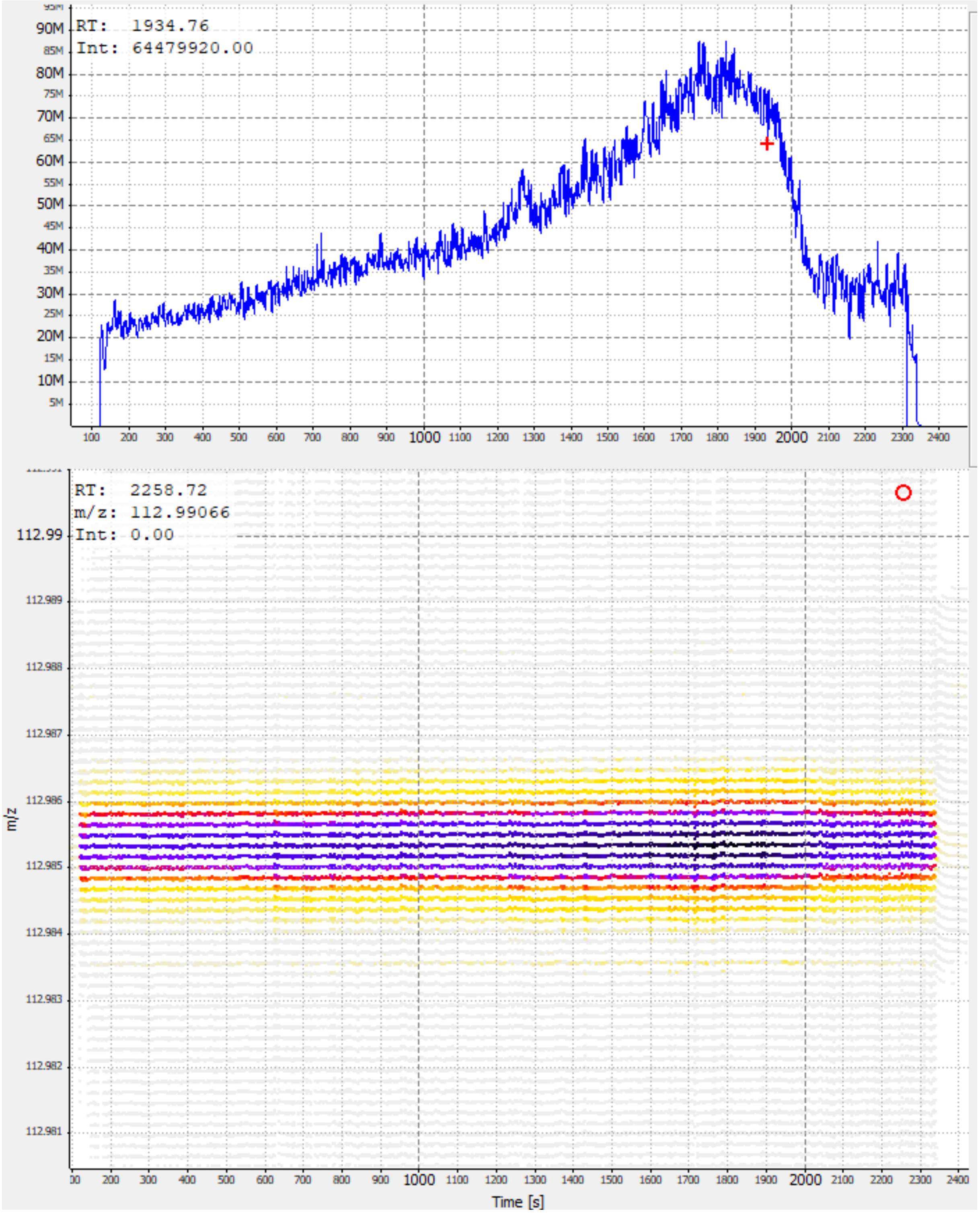

### Supplementary Figure 2

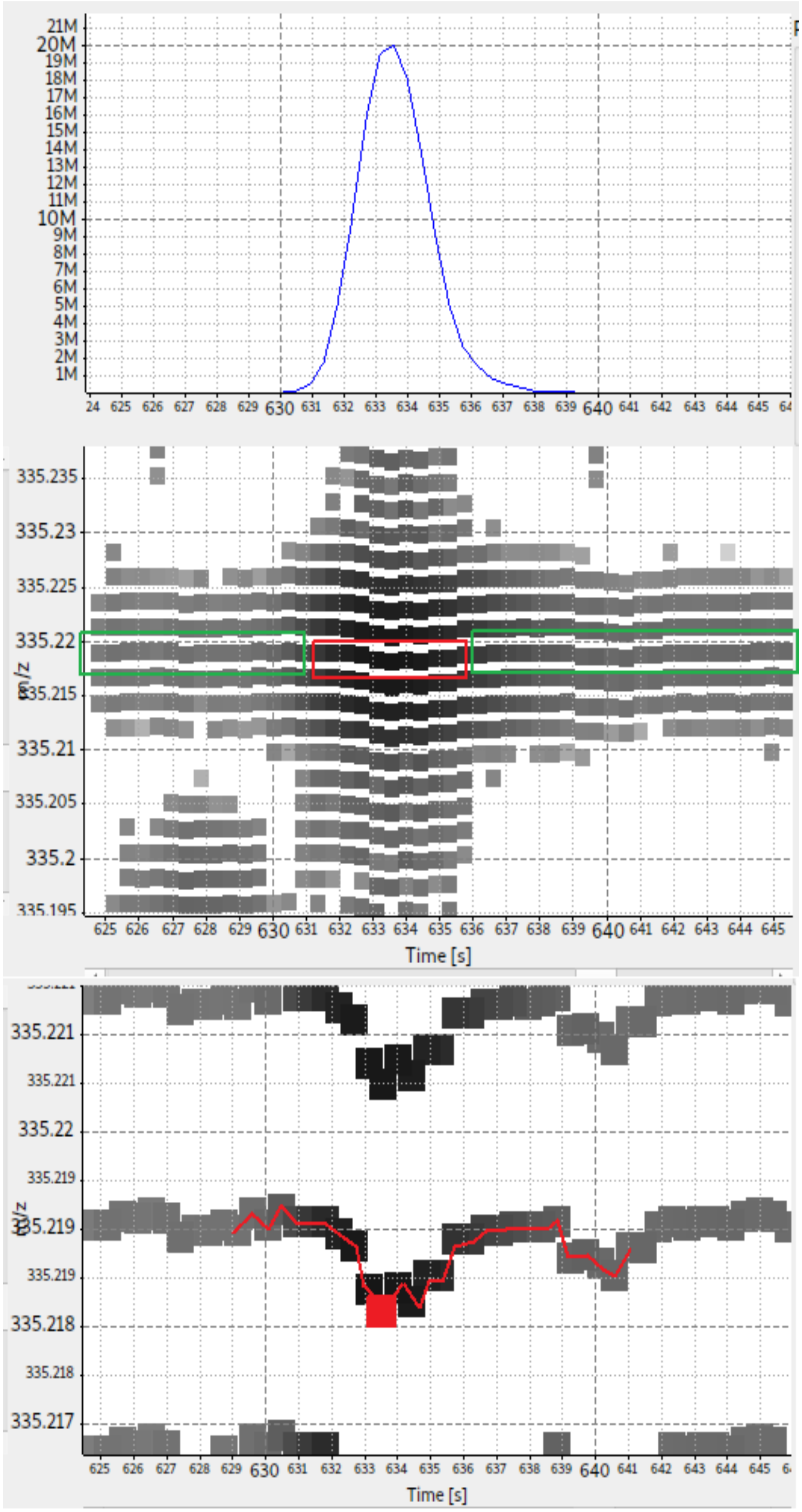

Highly abundant chromatographic peaks sometimes have shifts in the *m/z* values, which correlate with their chromatographic peak (top panel). These shifts typically are stronger in the peaks' centers where the chromatographic peaks have their highest abundances (red box in the middle panel) and less pronounced at the peaks' flanks where the chromatographic peaks are less abundant (green boxes in the middle panel). PeakBot accounts for such shifts and drifts by calculating the EIC adaptively. This is done by starting from the most abundant signal of the chromatographic peak (its apex, red box in bottom panel). Then the left and right neighboring signals with the most similar *m/z* value are chosen and the EIC is extended towards them. The two new *m/z* signals are derived from the previously chosen neighboring peaks and can be different for the left and right extension. This elongation step is repeated for a fixed number of scans (i.e., the retention time width of the standardized area). During each iteration of this extension phase the most similar *m/z* value is chosen as the new seed. Thereby even larger shifts can be accounted (red lines starting from the red box in bottom panel). In the depicted EIC example (red line), the difference between the highest and lowest *m/z* value was 0.001038 respectively 3.1 ppm. The illustrations were generated with TOPPView (Sturm and Kohlbacher 2009).

### Supplementary Figure 3

Overview of the training and validation matrices in the datasets not used for the development of PeakBot. Like the WheatEar dataset, which was used for the development of the PeakBot framework and that is presented in the main manuscript, each of these datasets has the 4 reference-sets T, V, iT, and iV (see the main manuscript for a description of these datasets). For each dataset a separate PeakBot model was trained with some 100 reference features and reference chromatograms. As can be seen, each model performs similarly well than the WheatEar dataset. Each dataset was used for 5 independent PeakBot CNN models (replicates; dots in the diagrams).

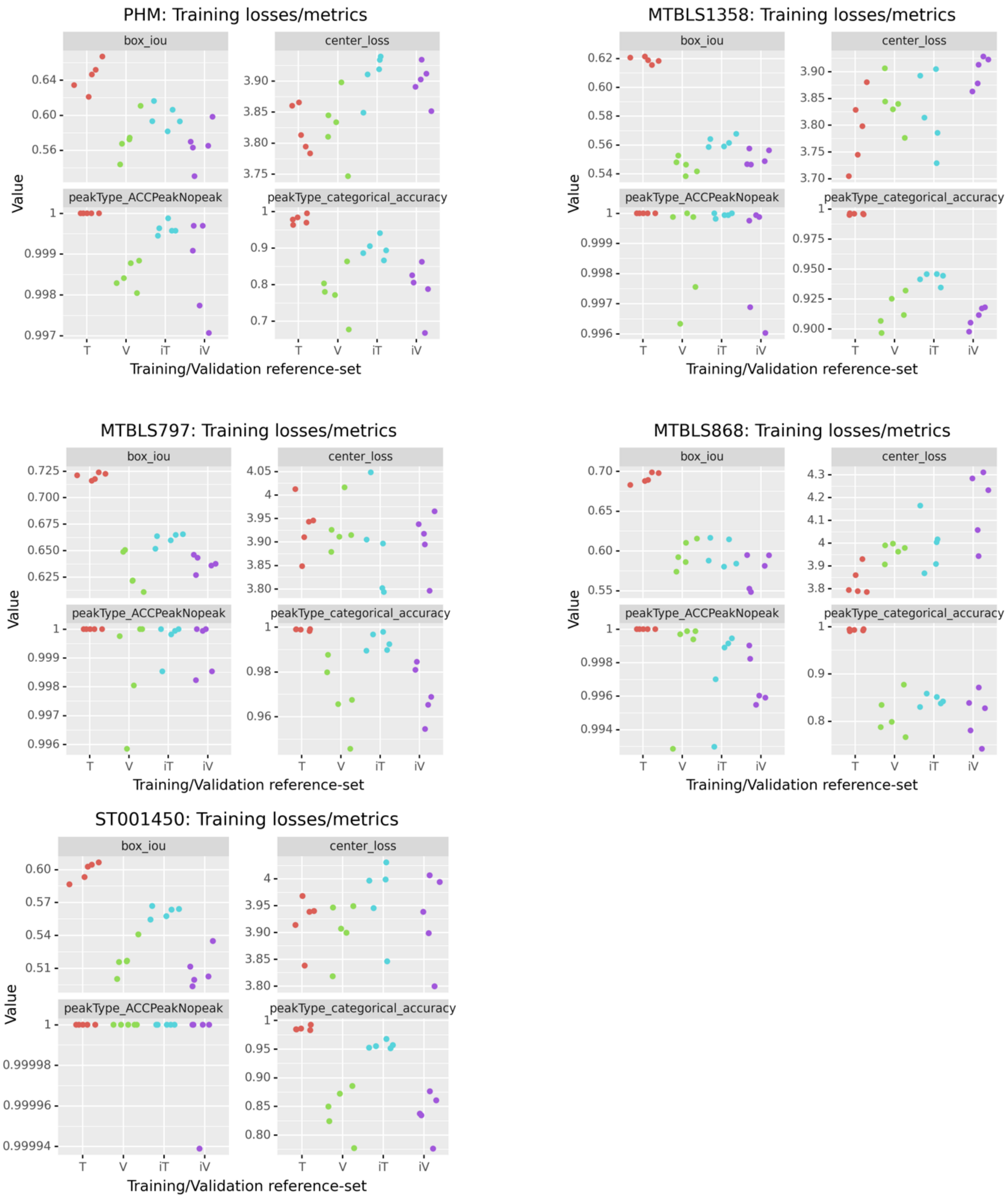

### Supplementary Figure 4

Illustration of the influence of using smaller or larger sized reference sets upon PeakBot's CNN model in respect to its loss/metrices. The different training/validation sets from the WheatEar dataset are shown. The center loss seems to be the least effected one, while the box and peakType are somewhat affected by a smaller number of reference features. The 6 peak-type classes (row PT_CatACC categorical accuracy) showed the largest degradation in accuracy. However, when the isomeric information (single peak, peak with earlier and/or later isomeric peak) are ignored, the true-positive and false-positive rates only change to a minor extend (rows PT_pFPR and PT_pTPR). Furthermore, the replicates for the same number of reference features indicate a randomized sub-selection of the total number of reference features (n = 2371) also showing the effect of different lists of ground-truth datasets. On the other hand, it is no surprise that when a small training dataset (10 to 50 reference features) are used to learn the characteristics of the chromatographic peaks, the model will overfit to the training data and present a high performance there while in the independent validation dataset (iV) the performance drops drastically. Thus, the performance is compared between the T and iV dataset to find the minimal number of 100 reference features (i.e., only minor differences in the metric values are observed).

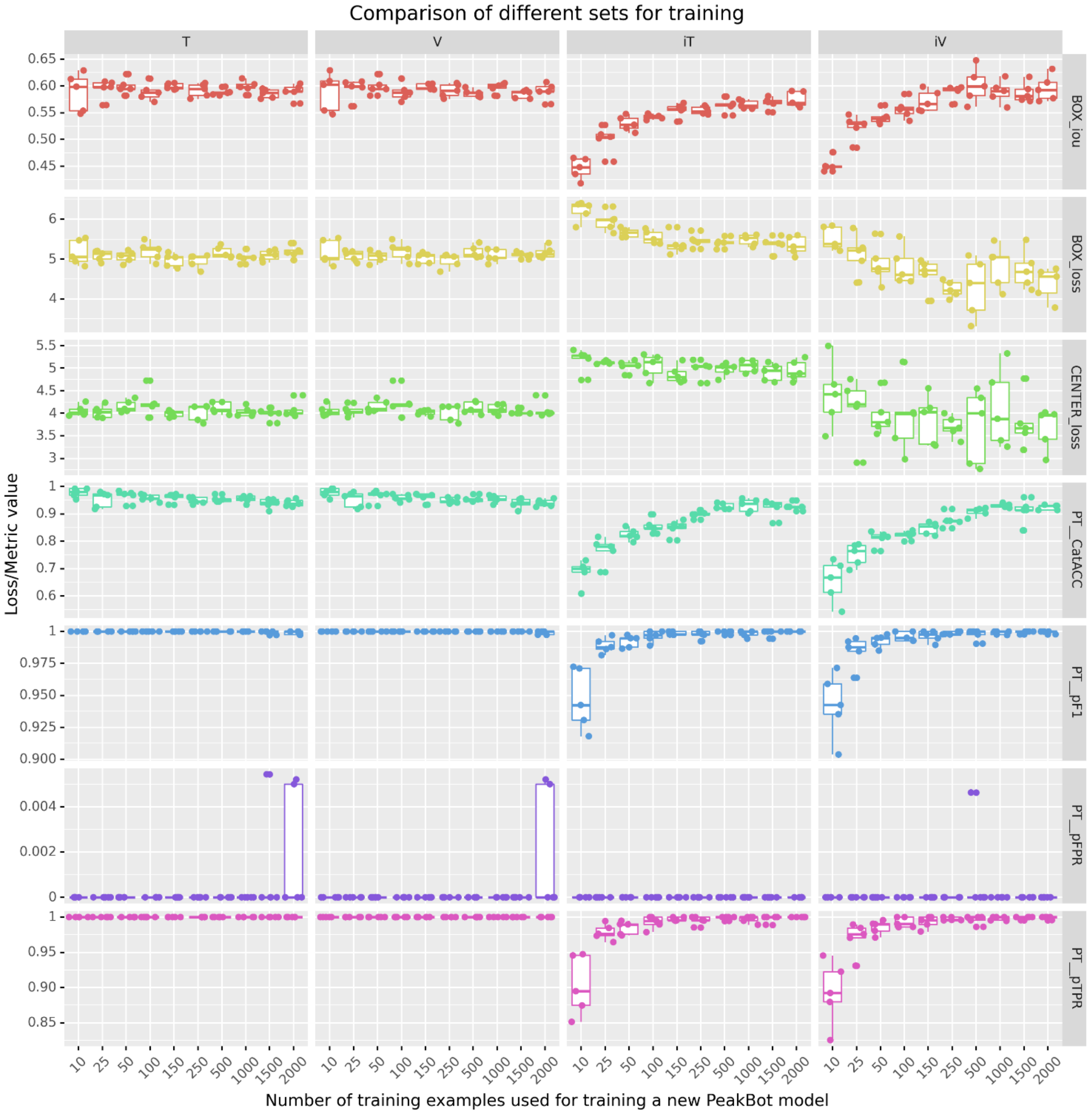

### Supplementary Figure 5

Illustration of the influence of using smaller or larger sized reference sets upon PeakBot's CNN model in respect to the total number of detected features in chromatograms. A lower number of reference features (10 to 50) drastically reduces the performance of detecting new chromatographic peaks not used for training in new samples.

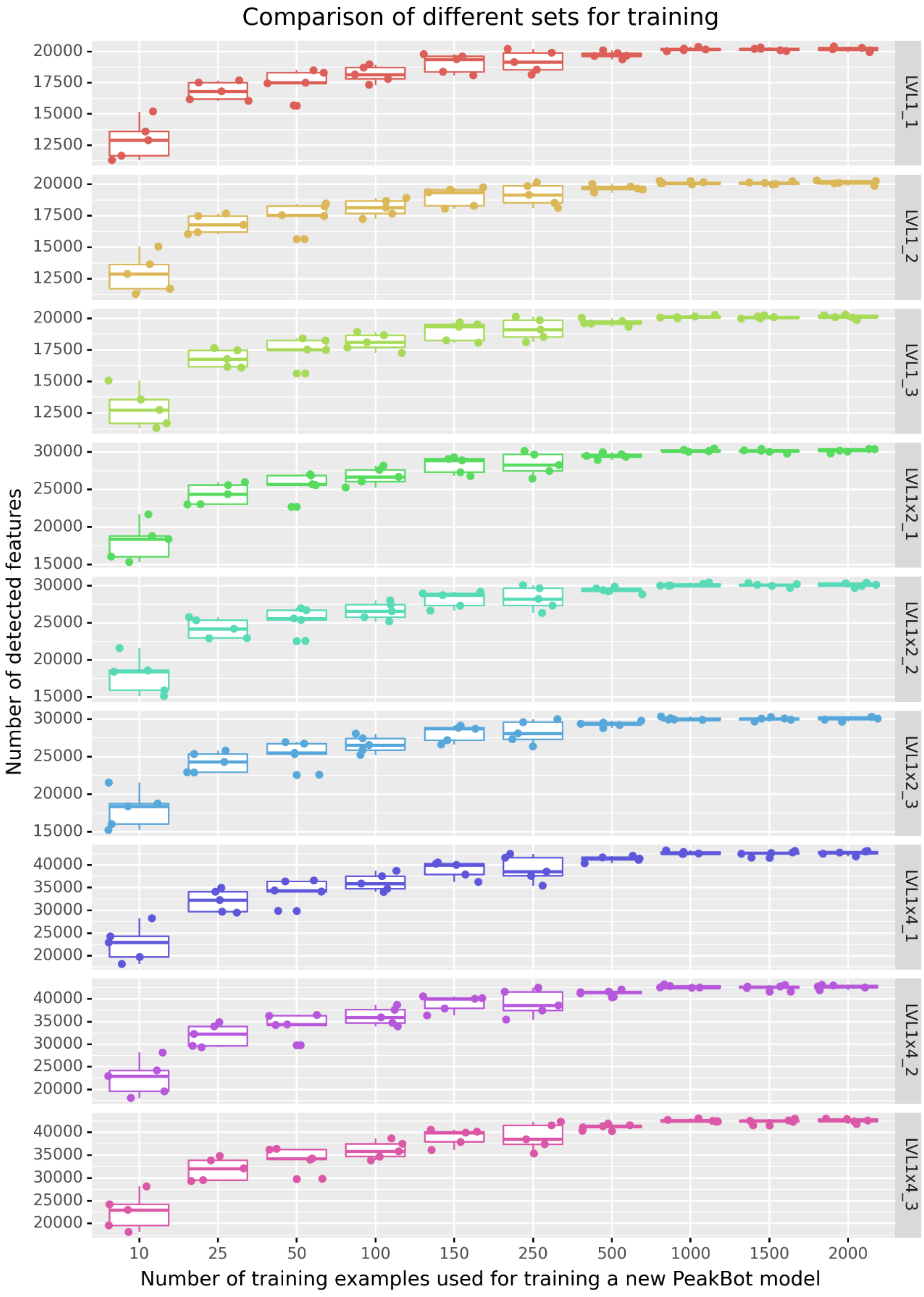

### Supplementary Figure 6

Comparison of the peak-picking results (number and overlap of the detected metabolic features) on the PHM dataset with the XCMS, MS-Dial, peakOnly, and PeakBot peak-picking tools. A high number of features (on average 64%), are successfully detected with all approaches. Moreover, on average an additional 13% of the features have not been detected with peakOnly. Also, on average less than 2% of features were exclusively detected with the two machine-learning assisted methods peakOnly and PeakBot. Each tool additionally found some features (on average less than 2 %) that were only detected with the respective tool.

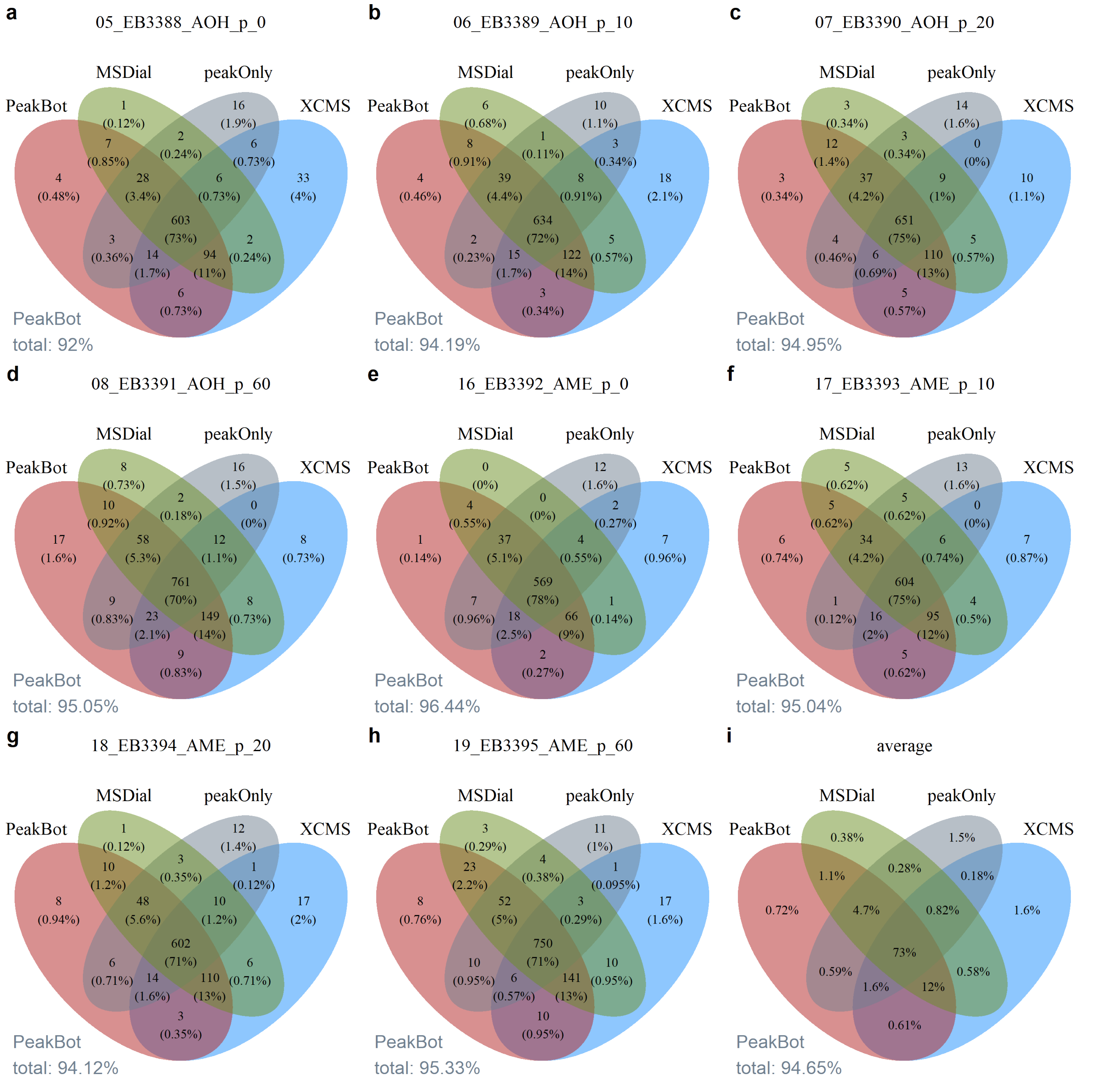

### Supplementary Figure 7

a Illustration of an MS/MS chromatogram (sample 823_sampleLvl1_Exp670_ddMSMS) obtained for the WheatEar sample with data dependent acquisition. Each black diamond indicates an MS/MS spectrum. As can be seen, several horizontal lines are present, which are background walls. Consequently, these MS/MS spectra have been "wasted" on non-informative scans. b shows another measurement (sample 823_sampleLvl1_Exp670_ddMSMS_wExList) of the sample shown in a, but this time the DDA was instructed to use an exclusion list, which contained the *m/z* and rt values of the background walls detected with PeakBot. As can be seen, many of these "wasted" MS/MS spectra have not been acquired. Note: The exclusion lists were only generated for the scans between retention time 2 and 36 minutes (120 and 2160 seconds) as outside of these scan ranges the chromatograms did not report reliable peaks.

a

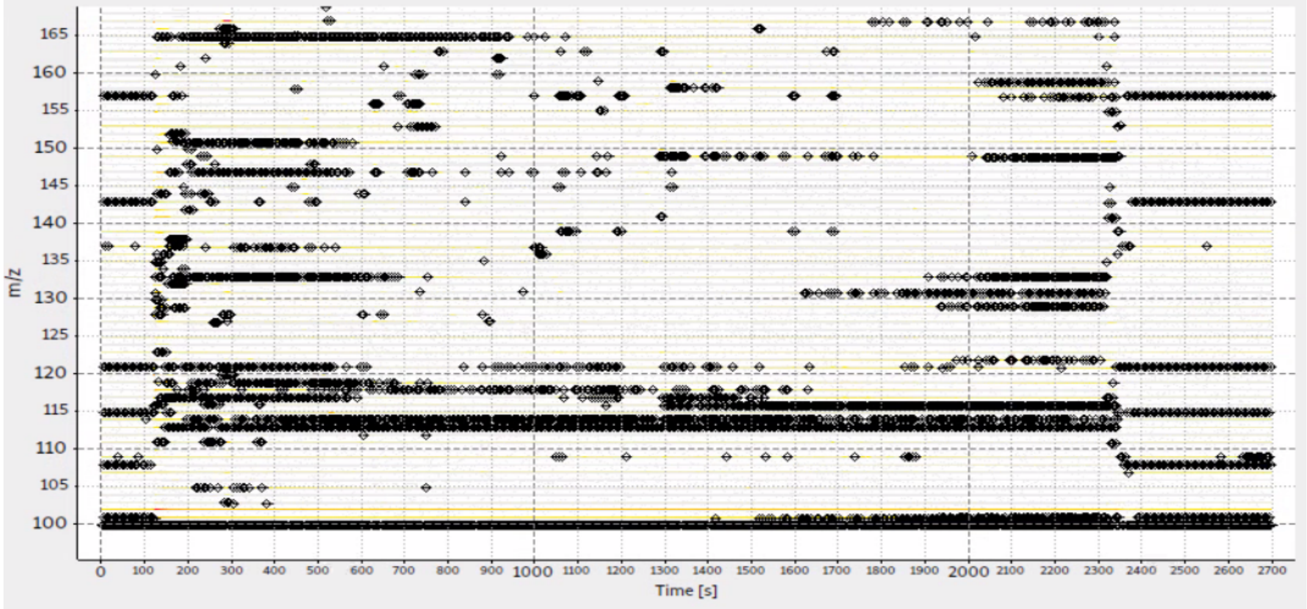

b

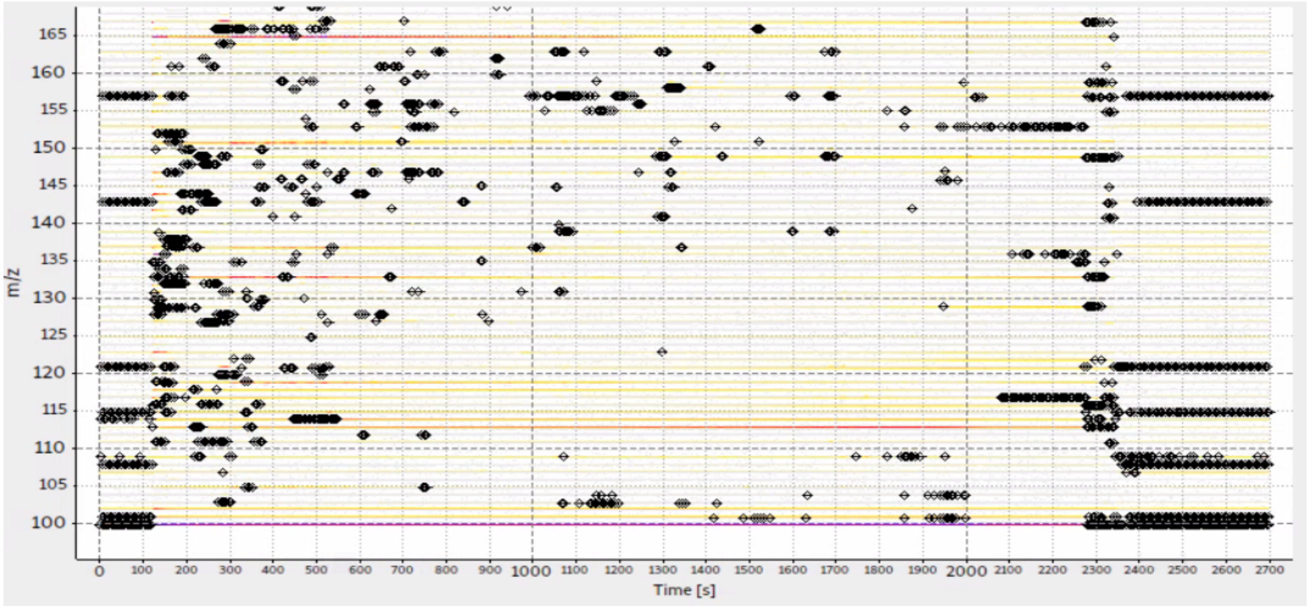

### Supplementary Table 1

Overview of local maxima, gradient descend peaks and PeakBot peak picking results for the MTBLS1358 dataset.

| Sample | Polarity | Local maxima | Gradient descend peaks | Chromatographic peaks | Backgrounds (e.g., walls) | Processing time (seconds) |
|---|---|---|---|---|---|---|
| HT_SOL1_LYS_010_pos_cm | + | 117,568 | 18,943 | 9,928 | 8,510 | 20.9 |
| HT_SOL1_SUP_025_pos_cm | + | 126,447 | 21,831 | 12,488 | 8,781 | 22.0 |
| HT_SOL2_LYS_014_pos_cm | + | 114,685 | 18,449 | 9,899 | 8,047 | 20.0 |
| HT_SOL2_SUP_029_pos_cm | + | 126,495 | 21,744 | 12,224 | 8,949 | 21.8 |
| HT_SOL3_LYS_018_pos_cm | + | 116,483 | 18,844 | 10,100 | 8,266 | 20.5 |
| HT_SOL3_LYS_033_pos_cm | + | 126,158 | 20,837 | 11,392 | 8,949 | 21.4 |

### Supplementary Table 2

Overview of number of MSMS spectra for true chromatographic peaks and for background walls with and without exclusion lists generated with PeakBot. [1]... excluding waste time of the chromatograms.

| Sample | Polarity | Exclusion list used | Total number of MSMS spectra [1] | MSMS spectra of chrom. peaks | Rate successful | Chrom. peaks with MSMS spectra |
|---|---|---|---|---|---|---|
| 823_SampleLvl1_Exp670_ddMSMS | - | No | 5,502 | 3,368 | 61.2 % | 1,356 |
| 823_sampleLvl1_Exp670_ddMSMS_wExList | - | Yes | 4,452 | 3,992 | 89.7 % | 1,605 |
| 823_SampleLvl1_Exp670_ddMSMS | + | No | 5,500 | 2,140 | 38.9 % | 855 |
| 823_sampleLvl1_Exp670_ddMSMS_wExList | + | Yes | 5,341 | 4,899 | 91.7 % | 1,810 |

### Supplementary Information 1

**Grouping of results from different chromatograms**

To track features from different samples after chromatographic peak picking with PeakBot, a nearest-neighbor algorithm is implemented in PeakBot. It can be executed multiple times with different parameters and samples. Each such iteration carries out the following steps:

1. All detected chromatographic peaks from all samples to be processed in this grouping step are first loaded into a long list.
2. The following steps are iterated until no feature is left without a group value:
    a. For each feature in this long list the nearest feature in both the retention-time and *m/z* dimension in a different sample is searched for. Here a user-defined retention time and *m/z* window is additionally tested and only two features within this window can be the nearest neighbors.
    b. The pair of two features closest together is taken and their group value is updated according to the following rules:
        i. If both features have not been assigned to a group, a new and unique group id is generated and both features are assigned to this group.
        ii. If either one these two closest features has already been assigned with a particular group, the other feature is updated with this group as well. Any feature that has the same group values as the one that needs to be updated will also be updated to the new value.
        iii. If for a certain feature no closest neighbor within the user-defined window is detected, then this feature is said to be alone and a -1 is used as the group value indicating that it cannot be grouped with another feature.
3. After the grouping has finished and all close-by features have received a group value or are alone, all such features are updated with the mean retention-time and *m/z* values of all its neighbors thereby accounting for small retention-time and *m/z* variations across the chromatograms.
4. The features with the updated group value are returned and this group information is used in the subsequent grouping iteration (if the user decided to use several such iterations).

### Supplementary Information 2

**Overview of wheat-ear samples (dataset WheatEar):**

Wheat samples were prepared as described in detail in (Doppler et al. 2019). The dataset consisted of three biological replicates of global metabolome labeling (GML, 1:1 mix of native and global $^{13}$C labeled plant material) samples of wheat ears harvested at flowering stage. LC-HRMS measurements were carried out with a Q-Exactive HF Orbitrap instrument coupled to a Vanquish UHPLC system (both Thermo Fisher Scientific, San Jose, CA, USA). A $C_{18}$-column (X-Bridge, 150 x 2.1 mm i.d., 3.5 µm particle size – Waters, Milford, MA, USA) was employed. $H_2O$ with 0.1% FA (v/v) (eluent A) and MeOH with 0.1% FA (v/v) (eluent B) were used for the chromatographic separation in a linear gradient that was 45 minutes long with a constant eluent flow of 250 µL min$^{-1}$ at 25°C (0 min: 10% B; 2 min: 10% B; 32 min: 100% B; 37 min: 100% B; 45 min: 10% B). Heated electrospray ionization (HESI) in polarity switching mode was utilized. MS1 data were acquired with fast-polarity-switching from *m/z* 100 to 1,000 with resolution set to 120,000 at *m/z* 200 in profile mode. Instrument performance was monitored by injecting a QC-reference standard mix consisting of 25 known metabolites. A detailed description of the LC-HRMS method including all chromatographic parameters can be found in (Sauerschnig et al. 2018).

The three biological replicates were measured with different injection volumes (2 µL for samples named “670_Sequence3_LVL1_x”, 4 µL (“670_Sequence3_LVL1x2_x”) and 8 µL (“670_Sequence3_LVL1x4_x”) with “_x” indicating the 3 replicate measurements) resulting in a total of nine files in this data set.

For testing the application of an exclusion list containing features classified as “walls” on the output of untargeted MS/MS measurements, 8µL of the wheat samples were injected and measured with the same chromatographic method as stated above. MS/MS fragment spectra were recorded with resolution 15,000 applying a top 5 data dependent acquisition method with dynamic exclusion of 3 seconds without (sample 823_sampleLvl1_Exp670_ddMSMS) and with (sample 823_sampleLvl1_Exp670_ddMSMS_wExList) using the generated exclusion list. The exclusion list for the latter sample was generated from the first sample 823_sampleLvl1_Exp670_ddMSMS. The DDA MS/MS acquisition settings were otherwise kept identically.

The acquired LC-HRMS data was converted to the mzXML format with msConvert (version 3.0.19228) from the ProteoWizard package (Chambers et al. 2012).

### Supplementary Information 3

**Overview of porcine hepatic microsome samples (dataset PHM):**

Porcine hepatic microsomes (PHM) were previously prepared in house, aliquoted and were stored at -80 °C until usage. The incubation mixes (3 mM NADPH, 1 mM ascorbic acid and 2 mg/mL microsomes in 200 mM Tris-HCl (pH 7.4)) were prepared on ice and after a 1 min pre-incubation (37 °C, 250-300 rpm) the toxin solutions were added (either alternariol (AOH) or its methyl ether (AME), 10 µM each) and incubated at 37 °C. After mixing (0 min) and after 10, 20, and 60 minutes incubation time a sub-sample was taken, mixed with three-parts of ice-cold extraction solution (acetonitrile/methanol, 1+1, v/v) and stored at -20 °C until further processing. Thereafter, the samples were centrifuged for 15 min at 17,000 g and 4 °C and transferred to HPLC-vials with microinserts. For LC-HRMS, 15 µL of each of the technical replicates (n = 3 to 7) were pooled and the pooled samples were analyzed both in positive and negative electrospray ionization mode. A Vanquish UHPLC system was coupled to a dual-pressure linear trap-quadrupole-orbitrap mass analyzer (LTQ Orbitrap Velos, both Thermo Fisher Scientific, Waltham, MA/USA) and was operated in full scan mode from *m/z* 100 to 1,000. Chromatographic separation was performed on a Supelco Ascentis Express C18 column (100 × 2.1 mm, 2.7 µm, Sigma Aldrich/Merck KGaA, Darmstadt, Germany) using a guard column of the same material. Eluent A consisted of aqueous ammonium acetate (5 mM, adjusted to pH 8.6 with a 25 % NH4OH) and eluent B was methanol. The flow rate was set to 0.4 mL/min and the following gradient resulting in a total run tome of 15.5 min was applied: The starting conditions of 10% B were hold for 1 min, followed by a linear increase to 100% at 11 min. Subsequently, the column was washed with 100% B for two minutes before returning quickly to the starting conditions for column equilibration.

The acquired LC-HRMS data was converted to the mzXML format with msConvert (version 3.0.19228) from the ProteoWizard package (Chambers et al. 2012). Only the positive ionization mode was used.

### Supplementary Information 4

**Overview of other datasets used for evaluating PeakBot's performance.**

Three publicly available datasets have been downloaded from the MetaboLights repository (https://www.ebi.ac.uk/metabolights). These datasets have the identifiers **MTBLS1358**, **MTBLS868**, and **MTBLS797**.

**MTBLS1358** is a study by Flasch and colleagues and investigated the metabolism of the mycotoxin deoxynivalenol in mammalian cell cultures (liver carcinoma cell line HepG2 and a model for colon carcinoma (HT29)). Both the medium of the cells and their intracellular matrix were analyzed with a Q-Exactive HF quadrupole-Orbitrap mass spectrometer. For further details about the biological study and analytical LC-HRMS measurements the interested reader is referred to the article (Flasch et al. 2020).

**MTBLS868** is a study by Mbekeani and colleagues by mining a fungal extract library for bioactive compounds against *Leishmania Mexicana*, a protozoan parasite that causes the cutaneous form of leishmaniasis. The authors analyzed their fungal extract library with a Q-Exactive Orbitrap mass spectrometer. Further details about the biological experiments and analytical LC-HRMS measurements are detailed in (Mbekeani et al. 2019).

**MTBLS797** is a study by Raheem and colleagues of extracts from different parts (bulb, leaf, scape and flower) of *Hyacinthoides non-scripta* flowers to analyze the metabolic compounds of the plant with molecular networking. The authors analyzed the plant extracts with an Exactive Orbitrap mass spectrometer. For further details about the biological study and analytical LC-HRMS measurements the interested reader is referred to the article (Raheem et al. 2019).

Another publicly available dataset has been downloaded from the Metabolomics Workbench repository (https://www.metabolomicsworkbench.org). Its identifier is ST001450.

**ST001450** is a study by Zhang and colleagues. A sample of human serum has been repeatedly measured with HILIC separation and a high-resolution LC-QTOF-MS instrument in the profile mode. For further details about the biological/technical aspects of the study and analytical LC-HRMS measurements the interested reader is referred to the article (Zhang, Dong, and Raftery 2020).

### Supplementary Information 5

**Comparison of XCMS, MSDial and PeakBot in respect to the overlap of commonly detected features.**

The samples from the PHM dataset were processed with XCMS (version 3.16.1), MS-Dial (version 4.80) and PeakBot. Only the peak-detection step was carried out with the following parameters for the tools:

**XCMS**: algorithm: centwave, peakwidth 2-5 seconds, 5 ppm mass deviation, signal to noise ratio threshold 5, integrate 1, *m/z* center function: wMeanApex3, mzdiff -0.0025, prefilter 1 x 1E5, noise 5E4, fit Gaussian peak. The centroided samples were used for this. For this, the vendor-centroiding algorithm of Thermo Fisher was used and the samples were converted to the mzXML format with msConvert from the ProteoWizard package.

**MS-Dial**: A project with LC separation and HRMS data obtained with Electrospray-Ionization was setup. Peak-detection parameters were 0.005 Da mass tolerance in MS1, Retention time begin and end 2 to 12.5 minutes, 0 to 2,000 *m/z* range, minimum peak height of 8E4, 0.005 Da mass slice window, linear weighted moving average smoothing method with a smoothing level of 3 scans, minimum peak width of 3 scans, no mass list exclusion. For this, the vendor-centroiding algorithm of Thermo fisher was used and the samples were converted to the mzML format with msConvert from the ProteoWizard package.

**peakOnly**: The raw data was converted to centroid mzML files and processed on a per-file basis with the available, pre-trained machine learning model. For this, the vendor-centroiding algorithm of Thermo Fisher was used and the samples were converted to the mzML format with msConvert from the ProteoWizard package.

**PeakBot**: A PeakBot-CNN model trained on the PHM dataset was used for detecting chromatographic peaks with PeakBot. The minimum intensity threshold was set to 1E5. For this, the samples were converted to the mzXML format with msConvert from the ProteoWizard package.

To find the features detected with the different approaches, all feature lists were compared. A maximum retention time and *m/z* deviation of 10 seconds respectively 10 ppm were allowed. If two or more features different approaches were similar (i.e., within the allowed rt and *m/z* deviation), a feature was said to be detected by all these peak picking approaches.

It must be noted here that a comparison of different software tools is challenging and that only few parameters can be transferred from one method to the other. Even worse, in many instances while parameter names are similar, they are being used by the internal algorithms differently. For example, in MS-Dial the minimum peak height parameter is considered only after baseline-subtraction (according to the developers' explanation available at http://prime.psc.riken.jp/compms/msdial/download/mathematics/MS-DIAL%20FAQ-vs2.pdf on page 10). On the other hand, both XCMS and PeakBot use this parameter differently and as an initial check for further consideration of signals to be used as ROIs or local maxima.

As a result of this, a lower peak height of 6E4 was used for MS-Dial here in this comparison. This reduced peak parameter allowed to find many peaks that would have been missed in the disfavor of MS-Dial.

Furthermore, for each detected feature with XCMS, MS-Dial and peakOnly, which operate on centroid mode LC-HRMS data, the profile-mode data was inspected if it contained at least one signal with an intensity of at least 1E5 thereby ensuring that the minimum intensity threshold of 1E5 is implemented in all four approaches identically. This was done as the centroiding step with msConvert sometimes slightly increases the abundance of the centroid signal in comparison to the profile mode data. This effect is expected as the centroiding step models the distribution of the *m/z* profile peak and thus the average value can have slightly higher intensities than the actually observed profile mode data.
